# Preparatory attention incorporates contextual expectations

**DOI:** 10.1101/2021.10.17.464696

**Authors:** Surya Gayet, Marius V. Peelen

**Affiliations:** Donders Institute for Brain, Cognition and Behaviour, Radboud University, 6525 HR, Nijmegen, The Netherlands; Helmholtz Institute, Department of Experimental Psychology, Utrecht University, 3584 CS, Utrecht, The Netherlands

**Author notes:** Corresponding author: *S. Gayet* –.

**Keywords:** Attentional selection, predictive processes, real-world vision, biased competition, visual search, scene perception, fMRI

## Abstract

Humans are remarkably proficient at finding objects within a complex visual world. Current theories of attentional selection propose that this ability is mediated by target-specific preparatory activity in visual cortex, biasing visual processing in favor of the target object. In real-world situations, however, the retinal image that any object will produce is unknown in advance; its size, for instance, varies dramatically with the object’s distance from the observer. Using fMRI, we show that preparatory activity is systematically modulated by expectations derived from scene context. Human participants searched for objects at different distances in scenes. Activity patterns in object-selective cortex during search preparation (while no objects were presented), resembled activity patterns evoked by viewing targets object in isolation. Crucially, this preparatory activity was modulated by distance, reflecting the predicted retinal image of the object at each distance. These findings reconcile current theories of attentional selection with the challenges of real-world vision.

**Highlights:**

- Visual cortex contains object-specific representations during search preparation.
- We demonstrate this for the first time during concurrent visual scene processing.
- Preparatory object representations are scaled to account for viewing distance.
- Preparatory biases reflect the predicted retinal image inferred from scene context.

**eTOC blurb:** Attentional selection is thought to be mediated by target-specific preparatory activity in visual cortex. Gayet and Peelen provide evidence that such preparatory biases incorporate contextual expectations about object appearance, reconciling attention theories with the challenges of naturalistic vision.

## Introduction

The vast majority of the sensory input entering through our eyes is irrelevant to our current behavioral goals. Consequently, the human visual system is equipped with means to favor behaviorally relevant visual input over irrelevant visual input. An influential idea is that attentional selection occurs by matching incoming visual input to attentional templates, so that objects matching (aspects of) the template receive prioritized processing ^[1]^ and attract spatial attention ^[2]^. At a neural level, such a template may be instantiated through the pre-activation of neural populations coding for the visual properties of the target object (e.g., its color, shape, size, or combination thereof). This preparatory activity serves to increase effective responsivity to visual input that comprises target-like properties, biasing competition in favor of task-relevant objects ^[3, 4, 5, 6, 7]^. The most direct evidence for such preparatory biases comes from monkey physiology and human neuroimaging studies that measured target-specific activity following search instructions but prior to stimulus onset. In this preparatory period, the firing rate of neurons tuned to target features increases within monkey inferotemporal cortex [8, 9], and stimulus-specific patterns of fMRI BOLD activity emerge throughout human visual cortex, ranging from primary visual cortex to inferotemporal cortex ^[10, 11, 12, 13, 14, 15, 16]^.

Despite these observations, there is ample reason to question whether the pre-activation of target-selective neurons in visual cortex is a viable selection mechanism for visual search in the real world. One key problem that is inherent to real-world visual search but is not accounted for by current models of attentional selection, is that a given object will produce a dramatically different image on the retinae depending on its location, which is unknown in advance. For instance, the color of the retinal image depends on the illumination on the object, its shape depends on the viewpoint, and – most critically – its size can vary by several orders of magnitude depending on the distance to the observer. Accordingly, the same object will evoke vastly different patterns of neural activity depending on its (yet unknown) location. In order to be an effective selection mechanism in the real world, preparatory activity could dynamically adjust to account for these situational dependencies. To account for the dependency between size and distance, for instance, this would entail pre-activating a smaller target object representation when searching far away, and a larger target object representation when searching for that same object nearby.

In the current study, we addressed this open question by measuring fMRI BOLD activity in human observers while they prepared to search for objects at different distances in indoor scene photographs. We considered three possible outcomes. First, preparatory activity may not play a role during naturalistic visual search at all, either because visual processing of a scene interferes with the concurrent maintenance of a target object representation ^[17, 18]^, or because the visual system cannot account for the location-dependency of the search target (as explained above). Second, preparatory activity may be situationally invariant ^[19]^. For example, preparatory activity may reflect a canonical (e.g., real-world) size of the target object, rather than its predicted – and variable – retinal image size ^[20, 21]^. Third, preparatory activity may incorporate the expected retinal size of the target object given the current viewing distance (i.e., more distant objects will produce a smaller retinal image). In line with this latter possibility, human observers occasionally fail to detect objects of inappropriate sizes given their location ^[22, 23]^.

We were able to distinguish between these possibilities by comparing the patterns of brain activity measured while observers were searching for an object at a given distance in a scene (Figure 1A; *search task runs*) to benchmark patterns of activity measured while observers were viewing those same objects in isolation in different sizes (Figure 1C; *training runs*). The experimental design of the search task had two critical assets. First, in order to ensure that activity measured during search only reflected preparatory activity, we analyzed data from a subset of trials in which observers prepared to search for the cued object, but the scene remained devoid of objects, and no actual search task was performed (Figure 1A, top). Second, we included different types of scene layouts (Figure 1B), to decouple viewing distance (i.e., ‘near’ and ‘far’) from the position of the cue (at the top or bottom of the image), thus requiring participants to extract the viewing distance from the scene itself rather than from the cue alone. The training runs with isolated objects were designed to retrieve benchmark activity patterns for both target objects (a melon and a box) in two different sizes (large and small). Object sizes in the training runs corresponded to the retinal image sizes that these objects would produce at the near and far distances of the search scene. These benchmark activity patterns allowed us to test directly whether preparatory activity (1) reflected a visual representation of the target object during search preparation, and (2) whether this visual representation was scaled to account for viewing distance during search. Taken together, the present experimental design captures a critical property of naturalistic visual search, incorporating contextual constraints on object appearance, while preserving full experimental control.

**Figure 1.**
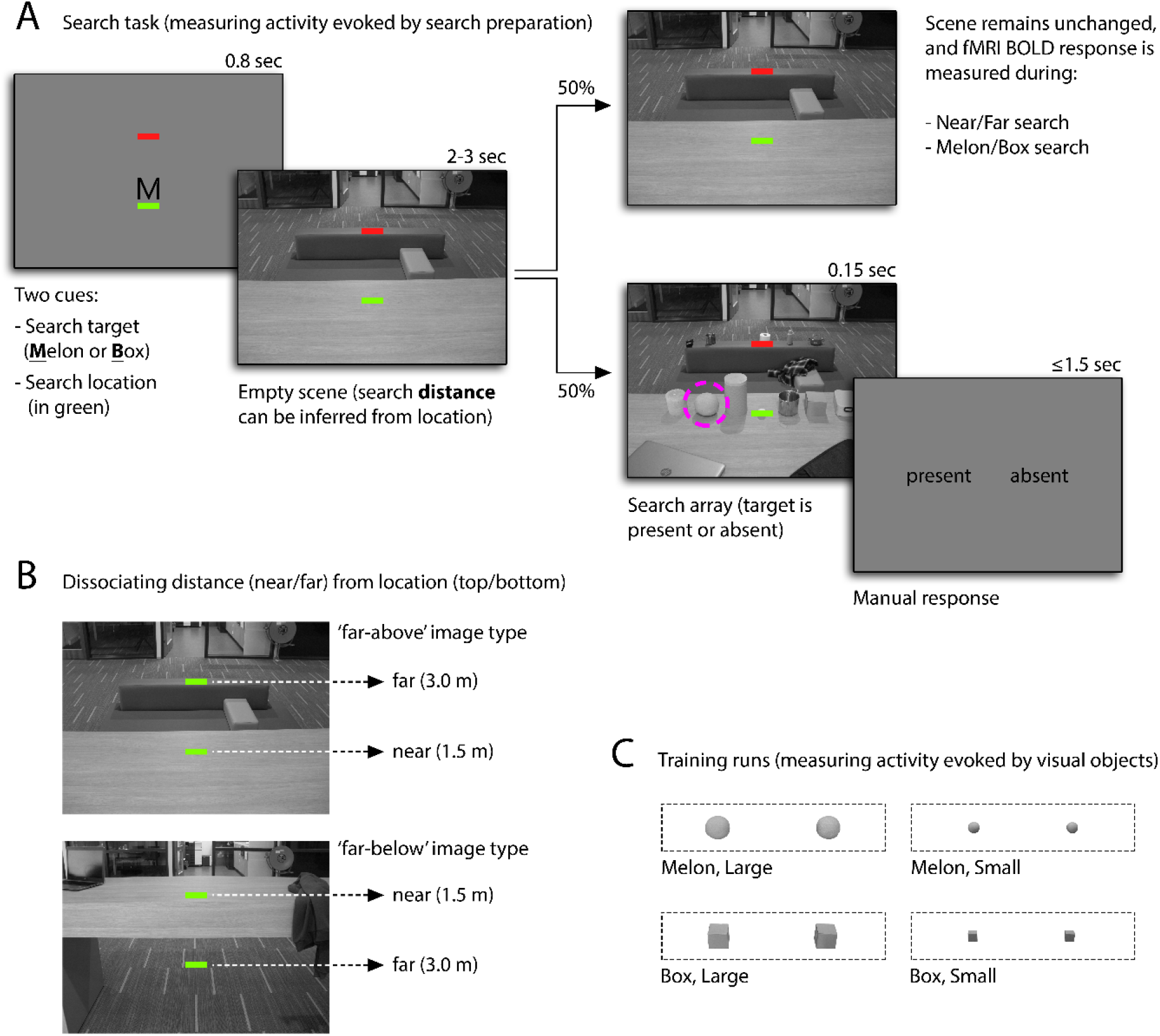
Schematic depiction of the experimental approach. (A) Search task. At the start of each trial, a letter (‘M’ or ‘B’) indicated which of two possible objects should be searched for (a melon or a box), and a green bar indicated the vertical location at which this target object could appear. After 800 ms, an indoor scene photograph was presented, which – together with the location cue (green bar) – enabled the observer to infer the real-world distance at which the target object should appear, thus predicting the retinal size of the target object. In half of the trials, after a varying delay (range 2-3 seconds) an array of 14 to 20 objects was briefly presented (150 ms), followed by a noise mask (100 ms; not displayed), and participants were required to report (within 1.5 seconds) whether the target object was present or absent (here: present; circled in purple for illustrational purposes). In the other half of the trials, the scene remained unchanged for the full delay period (i.e., no objects were presented), allowing for measuring fMRI activity evoked during search preparation (for near and far, melons and boxes). (B) The stimulus set included sixteen different scene families that categorically varied in their spatial lay-out: half of the scene families had the ‘far’ location in the upper half of the screen, and the other half had the ‘far’ location in the lower half of the scene. This manipulation ensured that distance information could not be inferred from the location cue alone, but had to be extracted from the scene instead. (C) The goal of the training runs was to reveal benchmark patterns of fMRI activity evoked by visual presentation of isolated objects (large and small melons and boxes), to compare those to the patterns of fMRI activity evoked during search preparation (for near and far melons and boxes). The isolated objects in these training runs were cropped from the search scenes and presented in a mini-block design.

To preview our findings: preparatory activity patterns in object-selective cortex during search preparation reflected the identity of the target object, resembling activity patterns evoked by visual presentation of these objects. Crucially, this resemblance was specific to comparisons in which the expected size during search matched the retinal size during visual presentation. We conclude that during naturalistic visual search (1) visual representations of the target object are pre-activated in object-selective cortex, and (2) are scaled to account for viewing distance. These findings demonstrate how preparatory biases in visual cortex can support visual search in the real world.

## Results

The main goal of this study was to test whether target-specific object representations – instantiated in visual cortex during search preparation – are scaled to account for viewing distance. A sample of 24 participants took part in two two-hour fMRI sessions on different days. Participants were cued to search for a target object (a melon or a box) in the near or far plane of an indoor scene photograph (see Figure 1A). In half of the trials, an array of objects briefly appeared at the cued distance (150ms), and observers reported whether or not the cued target object was present. This behavioral task was purposefully challenging, but feasible. Participants were 74% correct on target present trials, and 60% correct on target absent trials, amounting to 67% accuracy on overall present-absent judgements, which was better than chance; *p* < .0005. All reported *p*-values reflect the probability of incorrectly rejecting the null hypothesis (i.e., Type-1 error rate), based on 2000-samples bootstrap tests across participants (see STAR methods).

The fMRI analyses reported below are based on the other half of the trials, in which no objects appeared, thus isolating activity evoked by search preparation. Shared patterns of fMRI responses in visual cortex evoked by viewing an object in separate training runs (Figure 1C) and by preparing to search for that same object in the search task, are taken as evidence that this object is represented in the preparatory activity.

We focused our analyses on two main regions of interest (ROIs): object-selective cortex (hereafter OSC; mostly corresponding to the lateral occipital complex) and early visual cortex (hereafter EVC; mostly corresponding to V1 and V2). This followed from two main considerations. First, previous work has shown object-specific preparatory activity in these two regions while observers prepared to search for objects in naturalistic scenes ^[16]^. Second, these two areas constitute key regions underlying the perception of object size ^[24, 25, 26, 27, 28, 29, 30]^, as influenced by scene context ^[31]^. These ROIs were functionally defined for each individual participant (see STAR methods).

### Preparatory activity contains a target-specific object representation

Before asking whether target-specific object representations are scaled to account for viewing distance, we needed to ascertain that such object representations were indeed instantiated in visual cortex during search preparation. This is not a given, considering that all studies to date that demonstrated target-specific preparatory activity did so while observers were staring at an empty screen, in anticipation of an upcoming search task. By contrast, in our study we measured preparatory activity while participants were actively processing the scene in which they were searching for an object, as would be the case in real life. In a first analysis, we tested whether search preparation for melons and boxes (i.e., following an ‘M’ or ‘B’ cue in the search task) evoked distinguishable patterns of activity (irrespective of size). A multivariate classification approach (see STAR methods) showed that this was the case in OSC (*p* = .001), but we found no such evidence in EVC (*p* = .259), as shown in Figure 2A. As such, OSC retained information about the upcoming target object during search preparation. It is possible, however, that our analysis approach captured lingering activity evoked by the cues (i.e., an ‘M’ or a ‘B’) rather than sustained visual-like representations of the target objects.

**Figure 2.**
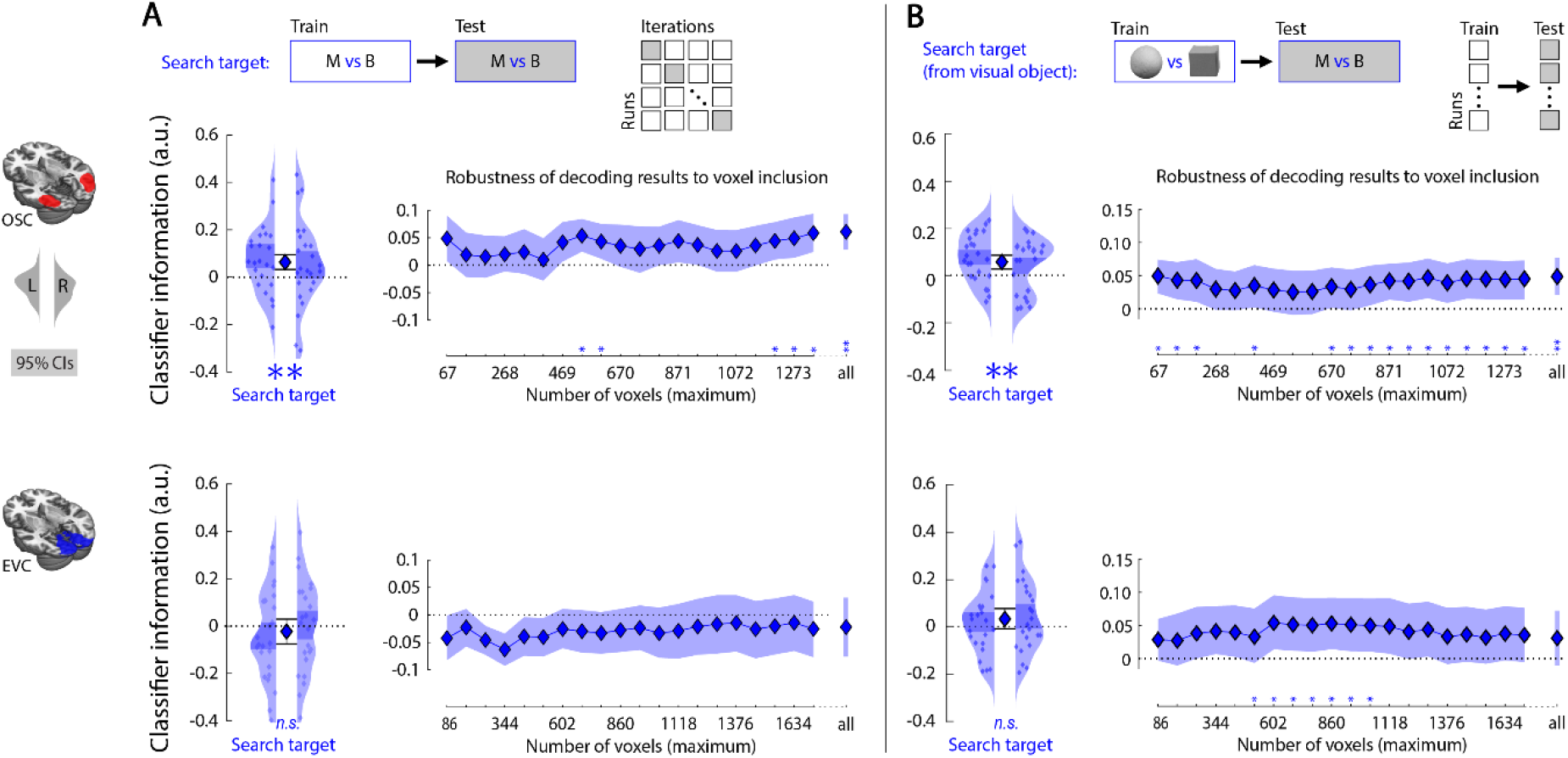
Analysis approach and results: decoding the target object during search preparation. (A) We first tested whether search preparation for melons versus boxes (i.e., following an “M” versus “B” cue in the search task) evoked distinguishable patterns of activity in OSC and EVC. This was the case in OSC, but we found no such evidence in EVC. Multivariate classification was achieved using a linear support vector machine (libsvm ^[32]^) on run-based beta-maps following a leave-one-run-out cross-validation procedure. To probe the robustness of the analyses conducted on the two default ROIs, we repeated these analyses for twenty different voxel inclusion thresholds (rightmost plots). Significance is reported after correction for multiple comparisons and threshold-free cluster enhancement ^[33]^ (see STAR methods). For reference, the default voxel inclusion threshold used in the main graph (on the left) is also included in these graphs, and labeled ‘all’. (B) Next, using a cross-classification approach, we tested whether a classifier trained on visually evoked activity in training runs (images of melons versus boxes) successfully cross-classified the objects that participants were cued to search for in the search task (melon versus box). This was the case in OSC. Small colored dots represent classifier information (derived from distance-to-bound) for individual participants, obtained separately from the left and right hemispheres (displayed within the left and right kernel-density plots, respectively). The central marker reflects the population mean, averaged across hemispheres. Error bars around the central markers, shaded areas within the kernel-density plots (on the left), and shaded area in the robustness plots (on the right), represent the bootstrapped 95% confidence intervals of the mean (2000 permutations). \**p* < .05, \*\**p* < .005, \*\*\**p* < .0005.

To distinguish between these possibilities, in a second analysis we tested whether activity patterns evoked during search preparation for the two target objects (while no object was actually presented) were qualitatively similar to those evoked by viewing the two target objects (i.e., images of melons and boxes presented in isolation in separate training runs, which evoked distinguishable activity patterns in both ROIs; *p* < .0005, for both tests). Again, for this analysis we lumped together large and small objects, and near and far search, to isolate the object-specific responses. Confirming our hypothesis, we observed above-chance object cross-classification in OSC (*p* = .004), but not in EVC (*p* = .098) (Figure 2B). These findings replicated across a wide range of voxel inclusion thresholds (Figure 2B, right-most panels), as well as using a reversed classification approach; a classifier trained to distinguish between search preparation for melons versus boxes successfully classified which object was visually presented in the training runs, based on activity patterns in OSC (Supplemental Information S1). Moreover, the generalization from visually evoked activity to preparatory activity was not driven by systematic differences in overall BOLD response to melon-compared to box-related conditions (Supplemental Information S2). Taken together, these findings show that activity patterns in OSC reflect a visual-like representation of the target object during search preparation. To our knowledge, this provides the first demonstration of preparatory object representations during concurrent scene processing.

### Preparatory object representations are distance-dependent

The main goal of this study was to find out whether preparatory object representations, as observed in OSC, are scaled to account for the current viewing distance. To test this, we first trained two different classifiers on visually evoked activity in the training runs: one classifier was trained to distinguish between activity evoked by viewing small images of melons versus boxes, and the other was trained to distinguish between large images of melons versus boxes. We could then compare the ability of both classifiers to distinguish whether participants were searching for a melon versus a box, separately for near search and far search. If object representations were rescaled during search preparation, the classifier trained to distinguish between *large* objects should perform best when cross-classifying target objects during *near* search and the classifier trained to distinguish between small objects should perform best when classifying target objects during *far* search. The data confirmed our hypothesis (Figure 3): when training on size-matching objects, preparatory activity in OSC yielded above chance cross-classification of the target object (*p* < .0005), but this was not the case when training on size-mismatching objects (*p* = .224; difference *p* = .017). Similarly, in EVC, preparatory activity yielded above chance cross-classification of the target object when training on size-matching objects, (*p* = .040), but not when training on size-mismatching objects (*p* = .244). However, cross-classification did not reliably differ between these two training regimes in EVC (*p* = 0.197). Confirming our finding in OSC, 19 out of 20 alternative voxel inclusion thresholds yielded an even more reliable difference-score than the default voxel inclusion threshold reported here (Figure 3, top-right panel). Moreover, the same conclusions were reached using a reversed analysis approach, in which classifiers were trained on activity evoked during search preparation and tested on activity evoked by visually presented objects (Supplemental Information S1). This size-specificity was not observed in face- and scene-selective regions of interest (Supplemental Information S3). Taken together, the present findings show that preparatory activity patterns in OSC reflect a representation of the search target that (1) resembles a visually evoked representation, and (2) takes into account the predicted retinal image size of the target object at the current viewing distance.

**Figure 3.**
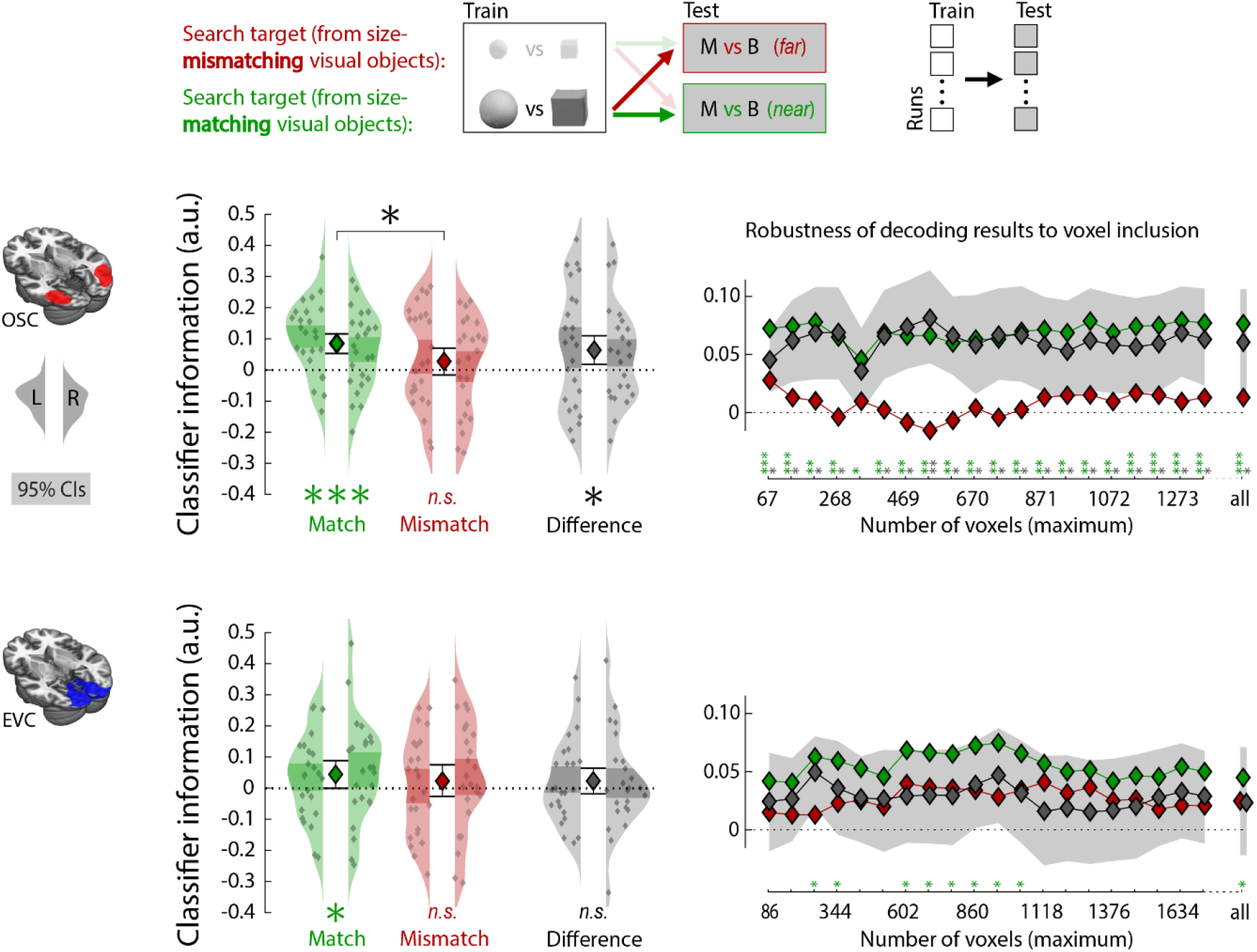
Analysis approach and results: distance-dependent object representations during search preparation. We tested whether cross-classification of target object (as shown in Figure 2B) improves when training a classifier on appropriately sized objects given the current viewing distance (e.g., large melons versus large boxes, when testing on *near* search) compared to inappropriately sized objects (e.g., large melons versus large boxes, when testing on *far* search). All analysis and data visualization approaches are identical to those described under Figure 2. For clarity, the robustness plots on the right only display confidence intervals for the critical difference score (size-matching minus size-mismatching object cross-classification). \**p* < .05, \*\**p* < .005, \*\*\**p* < .0005.

## General Discussion

We investigated whether target-specific preparatory activity is a viable mechanism for resolving competition between objects ^[3]^ in naturalistic conditions. To this end, we tested whether activity evoked during search preparation varies with viewing distance to account for the expected change in retinal image size of the target object. We observed that preparatory activity patterns in object-selective cortex contained information about the identity of the target object, scaled to match the predicted retinal size of the object at the current viewing distance. This preparatory activity could aid visual selection, by comparing the predicted target object representation to the actual visual input, thus prioritizing processing of the target object (e.g., template-based visual search ^[1, 2]^). The present findings demonstrate how preparatory activity can support attentional selection in the real world, where the visual input associated with an object depends on scene context: during search preparation, object-selective cortex contains a representation of the target object that flexibly adjusts to match the predicted retinal image that the target object produces at the current search location.

In addition to our main finding - that preparatory activity scales with viewing distance - the finding of object-specific preparatory activity (melon versus box) during naturalistic search constitutes an important generalization of previous work. Existing evidence for preparatory activity comes from studies in which observers searched for simple targets (e.g., vertical red bars) amongst distractors (e.g., red and green bars) on a uniform background ^[6]^. While natural scenes are arguably much richer than artificial displays (e.g., more clutter, occlusion), visual search in natural scenes is surprisingly efficient ^[34]^, and might therefore rely on different mechanisms than visual search in artificial displays ^[35, 36, 37]^. Thus far, very few studies have attempted to measure preparatory activity during naturalistic visual search (we are only aware of ^[16]^). Perhaps more importantly, all existing studies investigating preparatory activity recorded brain activity while participants stared at an empty screen in anticipation of the upcoming search task (i.e., following a cue that announced the target object). In real-world visual search, however, the outside world is not switched off during search preparation. Rather, observers concurrently process the visual scene in which they are searching for a target object. Active processing of a visual scene might interfere with the concurrent maintenance of a target representation in visual cortex ^[17, 18, 38]^, forcing observers to rely on more abstract parietal or prefrontal storage mechanisms ^[39, 40]^ (but see ^[41, 42]^). In the current study, a scene was presented during search preparation, and our data provides evidence that observers actively processed the scene as they successfully extracted distance information (as evidenced by the distance-dependency of object classification, depicted in Figure 2B). As such, the current study provides an important demonstration of target-selective preparatory activity during concurrent scene processing, showing that preparatory activity is a viable mechanism for real-world attentional selection.

It is well known that expectations modulate visual processing (see ^[43]^ for a review), causing expected visual objects to evoke a reduced but sharpened neural response ^[44, 45]^. Much like preparatory activity for explicit search targets, the expectation of an upcoming object suffices for evoking object-specific activity patterns in visual cortex ^[46, 47]^. In the present work, we show that observers used scene context to generate expectations about the appearance of an object. Specifically, knowing the distance at which an object was expected to appear allowed observers to generate strong predictions regarding the retinal size of the upcoming target object. Our results show that observers incorporated these predictions in their top-down attentional set, as reflected in combined distance- and object-specific preparatory activity.

We interpret the finding of distance-dependent object-specific preparatory activity as evidence that object representations are rescaled to match the current search distance. An additional preparatory mechanism that may play a role in real-world search is distance-based feature weighting; that is, the up- or down-weighting of object features based on their informativeness for the presence of the target object at a given distance. For instance, representing the texture of a cantaloupe melon might be more useful during near search (when texture is more visible) than during far search. Such a distance-based feature weighting mechanism could contribute to the current findings, as feature-specific preparatory activity (in the testing runs) may generalize to visually-evoked activity (in the training runs): classifiers trained to distinguish large melons from large boxes may rely more strongly on texture than classifiers trained to distinguish small melons from small boxes. Crucially, both the scaling and the differential-feature weighting interpretation support the same general conclusion that object-specific preparatory activity incorporates contextual expectations about target appearance. Future research is needed to isolate the features that are most important for distance-dependent updating of preparatory activity.

In the current study, participants were laying down with their head fixed, searching for pre-defined objects in two-dimensional gray-scale images following abstract cues. This setting differs in many ways from perception in the real world ^[48]^. The aim of the current study, however, was not to recreate real-world viewing conditions in the lab but, instead, to isolate one key aspect of real-world perception. In our study, viewing distance (as inferred from a photograph) determined the size with which a target object was represented during search preparation. This implies that participants were able to infer the viewing distance at which they were searching for a target object from our scene stimuli with sufficient precision using pictorial depth cues (e.g., perspective, blur, or comparisons to canonically-sized objects). During real-world vision, observers have many more cues at their disposal to obtain an even better estimate of viewing distance: binocular disparity, vergence, accommodation, and monocular movement parallax ^[49, 50, 51]^. Moreover, some of these real-world cues even provide information that is at odds with the available pictorial depth cues (e.g., vergence). For these reasons, extracting viewing distance is easier during real-world vision than in our experiment, presumably making distance-dependent preparatory activity an even more effective mechanism during real-world search than in the present experiment. At the same time, it remains unknown how the present paradigm in which observers were cued to search at two substantially distinct distances (1.5 and 3 meters) within a static display would generalize to attentional selection during dynamic visual input. Will preparatory object representations gradually change size as observers gaze across different distances, or will observers instead anticipate the viewing distance of the subsequent saccade endpoint to generate an object representation of the predicted size? Observers might continuously strike a balance between size-specific and size-invariant representations, depending on whether the gazing behavior moves parallel (e.g., looking at the road ahead) or transversal to the depth-axis (e.g., looking across the street). Having demonstrated that observers can utilize the inferred viewing distance to generate situation-specific preparatory biases, future work is needed to investigate how this mechanism is implemented in more dynamic settings.

Besides the retinal image size of an object, scene context provides various other predictions that could inform object processing ^[52, 53, 23]^. For example, the position of an object in a scene also predicts the shape of its retinal projection: a round clock on a wall to the left or right of the observer will be projected on the retinae as an ellipse. Behavioral work has shown that when observers searched for an elliptic target, their attention is captured by a canonically circular object (i.e., a coin) when it is rotated so as to produce an elliptic projection ^[54]^. This shows that, at the processing stage that is relevant to visual search, objects are (at least partly) represented in terms of their retinal projection, rather than their canonical real-world shape. Consequently, predicting the shape of the retinal projection of an object based on the angle of incidence could potentially benefit search. Object-selective cortex exhibits different degrees of invariance for such location-dependent factors as illumination, viewpoint, and size ^[26]^. As such, it remains unknown whether preparatory activity in visual cortex incorporates the angle of incidence similarly as it does for viewing distance. Nonetheless, the current work can be regarded as a proof-of-principle, showing that context-based predictions can be incorporated in preparatory activity, optimizing visual processing for goal-directed behavior in the real world.

There are at least two additional mechanisms that could work together with the preparatory scaling mechanism described here to support visual search at different distances. First, scene context could rapidly modulate the representations of target and distractor objects once these appear (or once they are fixated ^[55]^), perhaps even before competition is biased by preparatory activity. Behavioral studies have provided evidence for this notion, showing that spatial attention is captured by objects whose distance-inferred size matches the size of objects currently held in short-term memory ^[21]^, with such memory templates resembling preparatory activity ^[56, 57, 58]^. Second, preparatory activity can be instantiated at multiple levels of the visual hierarchy, from low-level features to high-level object categories ^[6]^. For some objects, it may thus be possible to pre-activate high-level representations that are invariant to size and viewpoint. In line with this, behavioral studies have shown that when observers searched for a highly-familiar object category (people or cars) in natural scenes, attention was automatically captured by outlines of a person (or car) regardless of their orientation ^[19]^. In sum, preparatory activity can comprise multiple features, each of which might be more or less diagnostic under different circumstances, and can be instantiated at different processing levels, allowing either for relatively early image-like selection, or for higher-level selection (e.g., view invariance). Considering human observers’ remarkable proficiency in detecting objects in naturalistic scenes ^[59, 60, 61, 62, 63, 64]^ it is probable that we flexibly switch between search mechanisms to optimally match situational demands.

## Conclusion

We proposed that for preparatory biases to work as a selection mechanism in naturalistic conditions, these would have to (1) operate when participants concurrently process visual information (i.e. looking at a scene while preparing to search), and (2) dynamically update as a function of the current search location (to account for viewing distance, illumination, angle of incidence, etc.). Here, we show for the first time that human observers generate visual-like representations of target objects in object-selective cortex while (1) actively processing the scene that they are searching, and (2) adapting these object representations as a function of viewing distance. By doing so, we demonstrate that preparatory biases in visual cortex are a viable mechanism for visual selection in the real-world.

## STAR Methods

### Resource availability

#### Lead contact

Further information and requests for resources should be directed to and will be fulfilled by the Lead Contact, Surya Gayet. This study did not generate new unique reagents.

#### Materials availability

All stimuli are publicly available at the Open Science Framework (OSF) project page associated with this study, which is listed in the Key Resources Table (https://osf.io/nqf6p/).

#### Data and code availability

- Behavioral data and processed fMRI data (beta-maps and regions of interest in MNI space) have been deposited at the OSF project listed in the Key Resources Table (https://osf.io/nqf6p/) and are publicly available as of the date of publication. Raw fMRI data in native subject space (activity maps and defaced structural scans) are deposited at the Donders Institution repository listed in the Key Resources Table (https://doi.org/10.34973/z3ym-9y81) and are publicly available as of the date of publication. Note that privacy regulations require one-time registration to access these data.
- Experiment scripts and analysis files (accompanied by comprehensive read-me files, and in-script annotations) have been deposited at the OSF project listed in the Key Resources Table (https://osf.io/nqf6p/) and are publicly available as of the date of publication.
- Any additional information required to reanalyze the data reported in this paper is available from the lead contact upon request.

### Experimental Model and Subject Detail

#### Participants

Participants were recruited through the Radboud university participant pool (SONA systems) and participated for monetary reward, after providing informed consent. The study was in accordance with the institutional guidelines of the local ethical committee (CMO region Arnhem-Nijmegen, The Netherlands, Protocol CMO2014/288).

A total of 24 participants (12 females, mean age = 24.1, SD = 5.2) took part in this study, and none were excluded. All participants completed two two-hour experimental sessions on separate days. The predetermined sample size of 24 followed from a trade-off between (A) the sample-size to achieve 80% power for obtaining an effect of medium size (N=34), and (B) our preference for having more within-subject power (i.e., two sessions instead of one), totaling to 48 experimental sessions. Maximizing the within-subject power was deemed necessary, because the hypothesized object-selective responses in preparatory activity were expected to yield small effect sizes, and key (decoding) analyses were performed within participants.

### Method Details

#### Apparatus

Participants viewed the stimuli through a mirror mounted on the head coil of the scanner. Stimuli were presented on a 1024 × 768 EIKI LC – XL100 projector (60 Hz refresh rate), back-projected onto a projection screen (Macada DAP diffuse KBA) attached to the back of the scanner bore. The effective viewing distance (eyes-mirror + mirror-screen) approximated 1440mm. Participants provided responses with the index fingers of the left and right hand, on a HHSC-2×4-C button box in each hand, connected to the serial port of the computer handling the stimulus presentation. All stimulus materials and experimental scripts described below can be found in the online repository, listed in the Key Resources Table.

#### General experimental procedure

Upon arrival at the scanner facilities, participants were guided through the three different tasks that they would perform inside the scanner (i.e., the search task, the training runs, and the functional localizer runs), as described below. Instructions were provided verbally, accompanied by a click-through demo and followed by a brief practice session on the experimenter’s laptop, to verify that participants understood the task instructions before going into the scanner. In the scanner, participants practiced the search task during the five-minute anatomical scan, while the experimenter monitored the behavioral responses. Participants performed a total of 24 functional runs. Each functional run started and ended with 15 seconds of fixation.

#### Experimental design & stimuli: Search task

The search task was designed to measure fMRI BOLD responses evoked during search preparation for different types of objects. Participants were instructed that they would search for one of two specific objects in scene photographs; a Cantaloupe melon and a small cube-shaped cardboard box (which was physically present during the instructions). On each trial, the letter ‘M’ or ‘B’ indicated whether they had to search for a melon or box, respectively. A green marker would indicate at what vertical location of the upcoming scene the target object could appear. After 1600ms, the scene appeared and, after a variable delay, an array of objects was briefly (150ms) presented within the scene. The array consisted of 14 to 20 objects, of which 12 to 18 distractor objects (drawn from of a pool of about 35 objects) such as a sweater, a laptop, a tea mug, a plant, and a watering can. The entire scene was then removed and backward masked (100ms), and participants were required to swiftly respond whether the target object was present or absent (within 1500ms), using button boxes in the left and right hand. After the response they would get feedback (correct, incorrect, or too slow), and the fixation dot changed color to announce the start of the next trial (1500ms inter-trial-interval). Participants were instructed that on half of the trials, no array of objects would appear within the scene, and hence no response would be required. Finally, it was stressed that participants were required to maintain fixation on the green marker throughout the trial, to ensure that they could detect the target object irrespective of its horizontal location (i.e., left or right of fixation). The two possible target locations always contained an object: either the target object, or a foil. The foil was either the non-target object (e.g., the melon when the box was cued) or an object that matched the shape (but not the size) of the target object (e.g., a football when the melon was cued on the far plane).

The study comprised sixteen search task runs, totaling to 512 trials per participant. Each search task run consisted of 32 trials, in an event-related design. Within each of these runs, five factors were counterbalanced within-participant and presented in random order. First, a green marker cued participants to fixate the upper part or the lower part of the upcoming scene (see Figure 1A). Second, one of two distinct types of scenes could be presented: scenes in which the far region was in the upper part of the image or the lower part of the image (see Figure 1C). Jointly, these two factors determined the search distance (i.e., near or far). Third, one of two possible object cues could be presented (i.e., the letter ‘M’ or ‘B’ indicating whether participants had to search for the presence of a melon or box respectively. Fourth, after a variable delay an array of objects appeared, or not. Fifth, on those trials in which an array of objects appeared, the cued target object could either be present in the array or not. One additional factor was counterbalanced with the five factors listed above across all eight search task runs of an experimental session, but not within each run: the scene background was chosen from one of eight scenes per scene type (sixteen scenes in total). Then a number of factors were (maximally) equated within runs, but could not be counterbalanced with the factors listed above. First, the type of distractor object at the non-target location could either be of a shape or a size that was similar to the cued target object. Second, the variable asynchrony between the scene onset and the onset of the search array could either be 2, 2.5 or 3 seconds (on trials in which no array was presented, the trial ended after the longest delay had elapsed). Finally, the mapping of the present/absent responses with the left/right button boxes was counterbalanced across participants. Participants never saw the same array of objects more than once.

The indoor scenes were photographed at sixteen different indoor locations within the Radboud University campus (Nijmegen, The Netherlands), using a digital camera on a tripod. A custom-made script was used to compute object positioning and camera angle, ensuring that the target objects would produce the same two retinal image sizes (large and small, corresponding to viewing distances of 1.5 and 3 meters), at the same eccentricity (left and right of fixation, in the near and far regions of the scene) in all sixteen different scene families. By doing so we ensured that the (predicted) sizes of the different target objects would be nearly identical within each of the two distance conditions (thus allowing for training binary classification algorithms). The images were turned grayscale, and any text appearing in the photographs was blurred using Photoshop 2017 (Adobe Inc.). The masks consisted of Pink (1/f filtered) noise, generated prior to each trial. The scene stimuli subtended 20 (width) by 14.4 degrees of visual angle (dva), and the target objects subtended about 0.9 by 0.9 dva (far) and 1.8 by 1.8 dva (near), with their inner edges positioned at an eccentricity of 3.2 dva from fixation. The upper and lower fixation positions were separated by 5.4 dva.

All sixteen different scenes (without objects), and all 512 unique search arrays used in the present experiment are publicly available via the Open Science Framework project page listed in the Key Resources Table.

#### Experimental design & stimuli: Model training

The purpose of the training runs was twofold: (1) retrieving benchmark activity patterns evoked by viewing isolated objects to train the multivariate models, and (2) identifying voxels that are visually responsive to the specific target stimuli used in our study (i.e., for defining the early visual cortex ROIs).

Participants were instructed to fixate the center of the screen, while pairs of objects were simultaneously presented to the left and right of fixation. Their task was to press a button whenever one of the two objects was 20% smaller or 20% larger than the remainder of the stimuli (i.e., an oddball detection task). The presentation of objects was subdivided into series (mini-blocks) comprising specific image categories (namely large and small melons and boxes), but this was irrelevant to the participants’ task.

Training runs consisted of a mini-block design, and the study comprised four training runs. Each run comprised 16 mini-blocks (four repetitions of four conditions, followed by a baseline fixation block). Each mini-block lasted 14.7 seconds and comprised 20 unique images (presented for 450 ms each, with a 250 ms blank in between). These images consisted of the target objects from the search tasks (i.e., melons and boxes) that were cropped from the scenes, equated in luminance, and presented on a uniform gray background. The objects were presented in pairs, one to the left and one to the right of fixation, at the exact same position (relative to fixation) as the target objects in the search task runs.

#### Experimental design & stimuli: Functional localizer

The purpose of the functional localizer runs was to identify object-selective voxels in individual participants. Participants were instructed to fixate the center of the screen, while different images were presented at fixation. Their task was to press a button whenever any image was presented twice in succession (i.e., a 1-back task). The presentation of images was subdivided into series (mini-blocks) comprising specific image categories (namely objects, scrambled objects, faces, and houses / landscapes), but this was irrelevant to the participants’ task.

The design of the localizer runs was identical to the design of the training runs, and the study comprised a total of four localizer runs. The stimuli and design of the localizer runs were based on (and nearly identical to) that of ^[65]^. In our set-up, the stimuli subtended 12 by 12 dva.

#### Acquisition of fMRI data

fMRI data were acquired on a 3T Magnetom PrismaFit MR Scanner (Siemens AG, Healthcare Sector, Erlangen, Germany) using a 32-channel head coil. A T2*-weighted gradient echo EPI sequence with 6x multiband acceleration factor was used for acquisition of functional data (TR 1s, TE 34ms, flip angle 60°, 2 mm isotropic voxels, 66 slices). For the search task, 295 images were acquired per run and 318 images were acquired per run for the training and localizer runs. A high-resolution T1-weighted anatomical scan was acquired at the start of each experimental session, using an MPRAGE sequence (TR 2.3 s, TE 3.03ms, flip angle: 8°, 1 mm isotropic voxels, 192 sagittal slices, FOV 256 mm).

#### Preprocessing of fMRI data

Data preprocessing was performed using SPM12. Preprocessing steps included field-map correction, two-step spatial realignment of the functional images, normalization to MNI 152 space (no down-sampling), and smoothing with a 3mm (FWHM) Gaussian filter. The two experimental sessions were independently warped into MNI space, and then combined.

#### Creating regions-of-interest: Object-selective cortex (OSC)

We ran a general linear model to model the responses evoked by viewing intact objects and scrambled objects in the localizer runs. Individual mini-blocks were modeled as boxcars and convolved with the canonical hemodynamic response function (HRF) provided in SPM12, and six motion parameters were included as nuisance regressors. Next, we computed a univariate contrast on the resulting run-based beta-maps to identify voxels that exhibited a significantly (*p*_uncorrected_ < .05) stronger response to intact objects than to scrambled objects. The ensuing collection of object-selective voxels for each participant was then intersected with a population-level functionally-defined object-selective mask (retrieved from ^[66]^), constraining the ROIs to lateral occipital regions, including the lateral occipital complex. The size of the OSC ROI varied across participants, with an average size of 1497 voxels (SD = 672; range = [181, 2693]) in the left hemisphere, and of 1173 voxels (SD = 512; range = [148, 2377]) in the right hemisphere.

To assess the robustness of our results, we created twenty additional ROIs for each participant, in which we differently constrained the maximum number of object-selective voxels. To this end, we sorted participants’ object-selective voxels within the group-level mask from most to least object-selective. Then, we created ROIs by keeping only the N most object-selective voxels within the mask, where “N” was a number increasing from zero to the median number of (significantly) object-selective voxels across all participants in twenty equidistant steps. Thus, this resulted in twenty additional OSC sub-ROIs of increasing size, and with increasingly liberal voxel inclusion.

All ROIs were initially constrained to a single hemisphere, allowing us to perform classification analyses within the left and right hemisphere separately (and averaging the results across hemispheres later).

#### Creating regions-of-interest: Early visual cortex (EVC)

We ran a general linear model to model the responses evoked by viewing target objects in the training runs (large and small melons and boxes). Individual mini-blocks were modeled as boxcars and convolved with the canonical HRF, and six motion parameters were included as nuisance regressors. Next, we computed a one-sample univariate contrast on the resulting run-based beta-maps to identify voxels that exhibited a significantly (p_uncorrected_ < .05) stronger response to all four object categories in the training runs (i.e., large and small melons and boxes) relative to the implicit baseline. The ensuing collection of visually responsive voxels for each participant was then intersected with an anatomical mask constituted of Brodmann’s Areas 17 and 18 ^[67]^, thus constraining the ROIs to a brain region mostly corresponding to primary and secondary visual cortex ^[68]^. The EVC ROI had an average size of 1713 voxels (SD = 604; range = [540, 2968]) in the left hemisphere, and of 1939 voxels (SD = 747, range = [690, 3244]) in the right hemisphere. Akin to the approach described above for the OSC ROI, we created twenty additional EVC sub-ROIs per participant that included an incremental number of visually responsive voxels. Again, separate ROIs were created for the left and right hemisphere.

### Quantification and Statistical analyses

#### Behavioral analyses

The search task comprised a total of 256 trials that required a behavioral response (i.e., in which an array of objects was presented), half of which were target-present trials, and half of which were target-absent trials. Trials that did require a response but in which no response was provided within the time limit were excluded from further behavioral analysis, so that the eventual analysis comprised an average of 126 (SD = 3.3) target-present trials, and 125.8 (SD = 3.4) target-absent trials per participant. Participants were 73.7% (SD = 9.7%) accurate on the remaining target-present trials (corresponding to a hit rate of 0.74), and 60.1% (SD = 13.3) accurate on target-absent trials (corresponding to a false-alarm rate of 0.40). To test for above-chance performance at the group level, we subtracted individual participants’ false alarm rates from their hit rates, to obtain a single performance metric for each participant. Then, on each of 2000 iterations, we drew N values from the resulting performance metrics, and computed the arithmetic mean (with N corresponding to the sample size of 24). To test for significant above chance-level performance, we computed the fraction of iterations (out of 2000) that yielded a positive average performance at the group level. If a positive value was observed on more than 95% of iterations, this was regarded as significant above chance performance, given an alpha level of 0.05.

Using the same general analysis approach, we observed no difference in behavioral performance between trials in which the target was a melon and trials in which the target was a box, *p* = 0.873. There was a small difference in behavioral performance between near targets (69% correct) compared to far targets (65% correct), *p* = 0.048, and between scenes in which the far plane was above the near plane (69% correct) compared to scenes in which the far plane was below the near plane (65% correct), *p* = 0.0016.

In the training runs, participants correctly reported 79% of size changes (i.e., hit rate) within our predefined response deadline of 1.5 seconds. Behavioral performance was slightly lower for melons (78% correct) than for boxes (80% correct), *p* = 0.0139, but did not differ between small and large objects, *p* = 0.810. It should be noted that no performance difference (e.g., between near-far or melon-box conditions) was shared between the search task runs and the training runs. It is therefore unlikely that successful cross-classification capitalized on differences in task difficulty.

#### General linear model (GLM) estimation

To model responses evoked by viewing target objects in the model training runs (large and small images of melons and boxes), we ran a general linear model on the data of each participant. Individual mini-blocks were modeled as boxcars and convolved with the canonical hemodynamic response function provided in SPM12. The GLM captured four conditions of interest, based on the factors ‘object size’ and ‘object type’: large images of melons, large images of boxes, small images of melons, and small images of boxes. To model responses evoked during search preparation in the search task, the search delays of individual trials (from scene onset to scene offset) were modeled as boxcars and convolved with the canonical HRF. Note that this included only the 50% of trials in which no array of objects appeared, and the scene thus remained unchanged and devoid of objects. For each participant, a single GLM was used to model all sixteen runs across two scanning sessions. This GLM captured the four conditions of interest based on the factors ‘search distance’ and ‘object cue’: near search for melons, near search for boxes, far search for melons, far search for boxes. In all GLMs, six motion parameters and one run-based regressor were included as nuisance regressors, and betas were estimated on a run-basis.

#### Multivariate pattern analyses

All multivariate classification analyses were performed with The Decoding Toolbox ^[69]^ using a linear support vector machine (hereafter SVM; libsvm ^[32]^). Classification analyses were performed on the run-based beta weights obtained from the GLMs (described in the previous paragraph), and were conducted within the left and right hemispheres separately. The results from both hemispheres were combined in the final step just prior to statistical testing. A leave-one-out cross-validation approach was used for the within run-type classification (i.e., within search task, or within model training runs), and a single-step cross-classification approach was used for the cross-classification analyses (e.g., training on visually evoked activity from the model training runs, and testing on activity evoked during search preparation in the search task runs).

For classification of object cue (melon versus box in the search task; Figure 2A), object type (melon versus box in the training runs), and cross-classification of object cue from object type (Figure 2B), all melon versus box maps were used, ignoring whether they were large or small, far or near. For the main analyses depicted in Figure 3 (cross-classification of object cue from size-matching and size-mismatching objects in the training runs) two separate classifiers were trained. First, a classifier was trained to distinguish between *small* melons and boxes, and tested on search preparation for melons versus boxes in the *far* plane (matching condition) and in the *near* plane (mismatching condition). Second, a classifier was trained to distinguish between large melons versus large boxes, and tested on search preparation for melons versus boxes on the *near* plane (matching condition) and in the *far* plane (mismatching conditions). The two matching conditions were then combined, and the two mismatching conditions were combined.

The number of training examples per classification analysis depended on the type of run used to train the classifier (there were 16 search task runs, and 4 model training runs), and the number of conditions included in the classification analysis (4 in total: large/near melon, small/far melon, large/near box, small/far box). To illustrate, we used 16 training examples per classification analysis to distinguish between visual presentation of melons versus boxes in the model training runs: 8 melon beta-maps and 8 box beta-maps (collapsed over size). Similarly, there were 64 test examples for the main classification analyses: 32 melon beta-maps and 32 box beta-maps.

To quantify the amount of information that is present in patterns of neural activity as retrieved by SVM classification (e.g., information about object type in OSC) we derived the following metric of classifier information from the distance-to-bound values *D*:

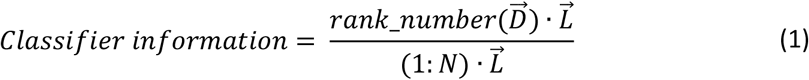

where N refers to the number of test examples for a given classification analysis (e.g., 64 for melon versus box classification in search task runs); D is an array of length N comprising the distance-to-bound values for each of the N test examples; and L is an array of length N containing the N labels (i.e., either −1 or 1) describing the correct classification category of the corresponding values in D (e.g., “-1” for “melon” and “1” for “box”). Put simply, the numerator of this equation reflects the amount of classifier information (positive for correct classification and negative for incorrect classification), and the denominator is a constant, which normalizes the classifier information metric to the range [-1, 1].

The sign of the distance-to-bound values reflects binary classification decision, with negative and positive values reflecting classifier decision “melon” and “box”, respectively. The magnitude of these values can be regarded as the classifier’s certainty of this classification decision. By computing the dot product of these distance-to-bound values (negative for “melon” and positive for “box”), with their associated correct test labels (“-1” for “melon” or “1” for “box”), the ensuing metric provides an increasingly positive value for increasingly correct classification and an increasingly negative value for increasingly incorrect classification. Instead of using the raw distance-to-bound values, we use their rank values (where the most negative value is ranked 1, and the most positive value is ranked N). This is important, because the hyperspaces obtained from different linear classifiers are not directly comparable (e.g., when averaging across hemispheres or conditions of non-interest, or comparing across conditions of interest and participants). Using this approach, we remain agnostic to the shape or space of the different hyperspaces, while still penalizing incorrect classifications more when they are very ‘certain’ (e.g., rank 1 or 64 out of 64) more than when they are very uncertain (e.g., rank 32 or 33 out of 64).

This metric of classifier information has two crucial advantages over traditional binary classification accuracy. First, continuous (i.e., distance-to-bound) measures are more sensitive than binary classification decisions ^[70]^. Second, this approach is inherently immune to classification biases that systematically favor one label over the other. Importantly, because our metric relies on rank order, it does share the key advantage of binary classification metrics: being agnostic as to the shape or range of the hyperplane of the linear classifier, thus allowing for comparison across multiple classifiers.

#### Significance testing

For each main ROI (e.g., OSC), and each condition of interest (e.g., classification of object type within-training runs) we performed bootstrap tests against chance, with 2000 samples (allowing for a lower bound of *p*_MIN_ < .0005). Specifically, on each iteration, we drew 24 samples with replacement from the 24 classifier information metrics (i.e., one for each participant), and computed the arithmetic mean. To test for significance, we then computed the fraction of iterations (out of 2000) that yielded above chance-level classifier information at the group level. If a positive value was observed on more than 95% of iterations, this was regarded as significant above chance-level classifier performance, given an alpha level of 0.05. Note that this approach entails a directional test, which followed from the strong prediction that classifier information should be either at chance or above chance but not below chance.

To evaluate the robustness of our results observed in our primary ROIs, we conducted each classification analysis in each generic ROI (EVC and OSC) an additional twenty times, in the twenty sub-ROIs of increasing size (see ROI description above). In all twenty additional analysis, a new SVM was trained and tested (following the same classification procedure described above) and classifier information metrics were computed as described above. In addition, we also computed classifier information metrics after pseudo-randomly permuting the correct test-labels (e.g., melon or box) against which the classifier outcomes were pitted. Importantly, the same (permuted) labels were used across all twenty sub-ROIs, to preserve inherent correlations between sub-ROIs. This procedure was repeated 2000 times to generate a null distribution that has the same variance and autocorrelations as the actual data, but should not carry any information about object type. Next, we applied threshold-free cluster enhancement ^[33]^ (TFCE) using the CoSMoMVPA toolbox ^[71]^. TFCE boosts belief in consecutive data-points with signal (i.e., representing cluster-like spatial support of individual data-points). TFCE-scores were computed for the observed data as well as the null data. The eventual test statistic, as reported in the figures, conveys how likely a given TFCE-score is (for each sub-ROI), given the maximum TFCE-score across all sub-ROIs in the null data, thus accounting for cumulative Type I error. Importantly, using this method, statistical significance establishes the existence of above-chance classification; it does not allow for making claims about the location or extent (i.e., sub-ROIs) of this effect ^[72]^.

### Key resources table

| REAGENT or RESOURCE | SOURCE | IDENTIFIER |
| --- | --- | --- |
| Deposited Data |  |  |
| Raw data (fMRI & behavior) | This study | https://doi.org/10.34973/z3ym-9y81 |
| Processed data (fMRI & behavior) | This study | https://osf.io/nqf6p/ |
| Stimulus material (training and search task runs) | This paper | https://osf.io/nqf6p/ |
| Stimulus material (functional localizer runs) | [65] | https://osf.io/nqf6p/ |
| Population-level regions-of-interest | [66] | https://osf.io/nqf6p/ |
| Software and Algorithms |  |  |
| MatLab (versions r2018a, r2021a) | Mathworks | http://www.mathworks.com/products/matlab/; RRID: SCR_001622 |
| SPM (version 12) | [73] | https://www.fil.ion.ucl.ac.uk/spm/software/spm12/; RRID: SCR_007037 |
| The Decoding Toolbox, TDT (version 3.98) | [69] | https://sites.google.com/site/tdtdecodingtoolbox/; RRID: SCR_017424 |
| CosmoMVPA (last git pull: 16-12-2020) | [71] | http://cosmomvpa.org/; RRID: SCR_014519 |
| Psychtoolbox (version 3.0.14) | [75, 75] | http://psychtoolbox.org/; RRID: SCR_002881 |
| Image Processing Toolbox (version r2018a) | Mathworks | https://nl.mathworks.com/products/image.html |
| Other |  |  |
| MatLab code for running experiment | This study | https://osf.io/nqf6p/ |
| MatLab code for fMRI data pre-processing, region-of-interest creation, analyses, and statistical testing | This study | https://osf.io/nqf6p/ |
| MatLab code for analysis of behavioral data | This study | https://osf.io/nqf6p/ |

## Author contributions

SG and MVP developed the study concept and study design. SG programmed the experimentation scripts, and developed the pre-processing and analysis scripts in consultation with MVP. SG tested the participants. SG drafted the manuscript and MVP provided critical revisions. Both authors approved the final version of the manuscript.

## Acknowledgments

This project received funding from the European Research Council under the European Union’s Horizon 2020 research and innovation program (Grant Agreement No. 725970, granted to Marius V. Peelen), and from the Netherlands Organisation for Scientific Research (Vl.Veni.191G.085, granted to Surya Gayet). The authors would like to thank Nicolò Trevisan for help with the stimulus creation and data collection, and Stefan van der Stigchel, Maëlle Lerebourg, and Genevieve Quek for helpful comments on earlier drafts of the manuscript.

## Diversity statement

Recent work in several fields of science has identified a bias in citation practices such that papers from women and other marginalized groups are under-cited relative to the number of such papers in the field ^[76, 77, 78, 79, 80, 81]^. We seek to proactively consider choosing references that reflect the diversity of the field in thought, form of contribution, gender, and other factors (but acknowledge that we likely failed to fully account for the bias against these underrepresented groups). We decided against reporting numerical metrics classifying the authors represented in our reference list (e.g., using databases that store the probability of a name being carried by a woman ^[80, 81]^), for two main reasons. First, reporting binary classification of gender marginalizes intersex, non-binary, and gender non-conforming people. Second, selectively reporting gender (or classifying along one or more other dimensions) marginalizes all other under-represented groups. With this statement, we want to (at least) raise awareness of this issue, and look forward to future work that could help us to better understand how to support equitable practices in science.

## Supplemental Figures

**Supplemental Figure S1.**
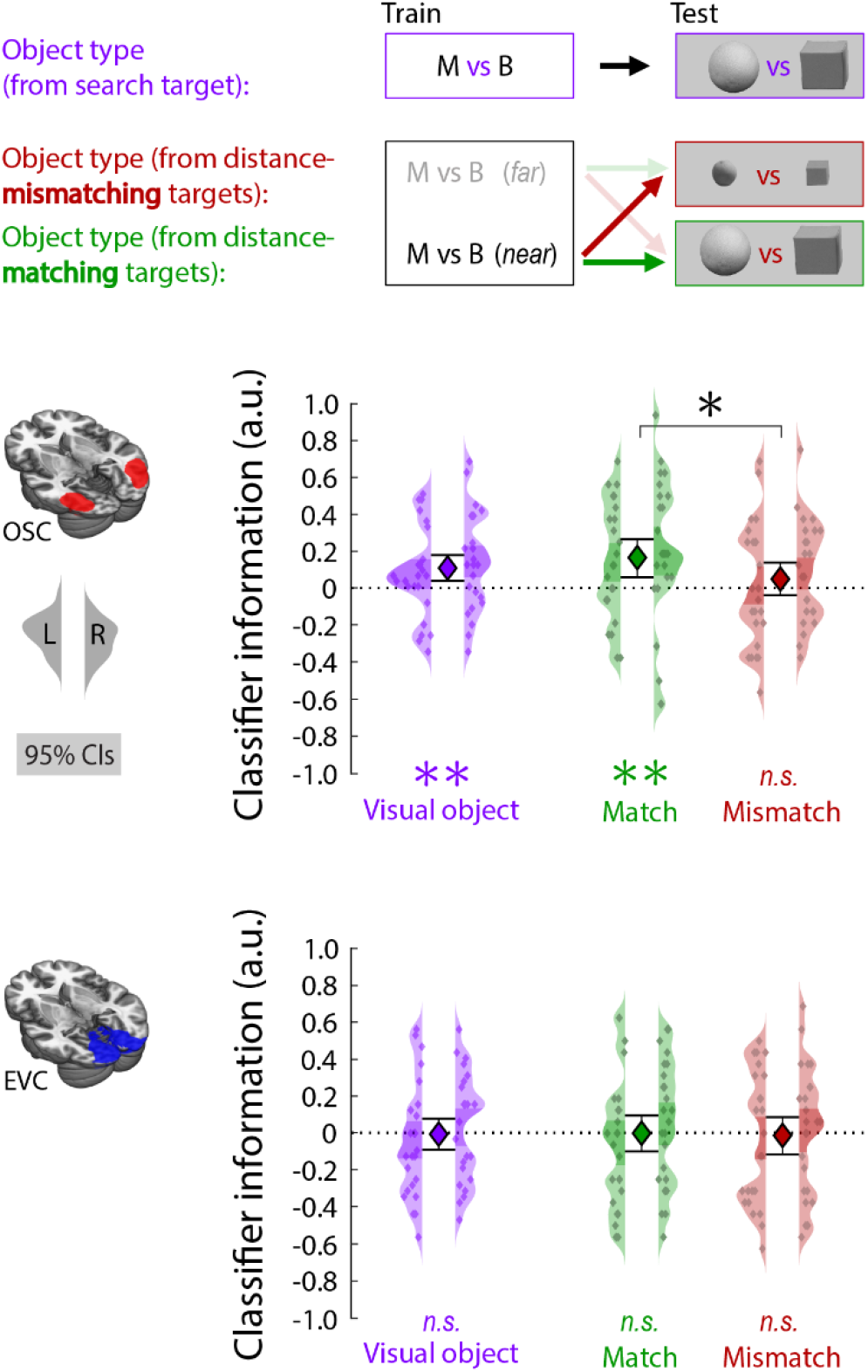
To corroborate the cross-classification results reported in the main manuscript, we replicated our main analyses after reversing the cross-classification direction which should lead to qualitatively similar results despite being based on completely different testing and training datasets ^[S1]^; although they might differ quantitatively ^[S2]^). This entailed training the classifier on activity evoked during search preparation (in the search task) and testing the classifier on activity evoked by viewing isolated visual objects (in the training runs). First, we tested whether a classifier trained to distinguish between melon versus box search (i.e., following an ‘M’ versus ‘B’ cue) could distinguish between visually presented melons versus boxes (in purple). Second, we tested whether this melon versus box cross-classification improved when training on the appropriate search distance (in green) compared to the inappropriate search distance (in red). All key findings reported in the main manuscript were replicated using this reversed cross-classification approach. Small colored dots represent classifier information (derived from distance-to-bound) for individual participants, obtained separately from the left and right hemispheres (displayed within the left and right kernel-density plots, respectively). The central markers reflect the population mean, averaged across hemispheres. Error bars around the central markers, and shaded areas within the kernel-density plots represent the bootstrapped 95% confidence intervals of the mean. \**p* < .05, \*\**p* < .005.

**Supplemental Figure S2.**
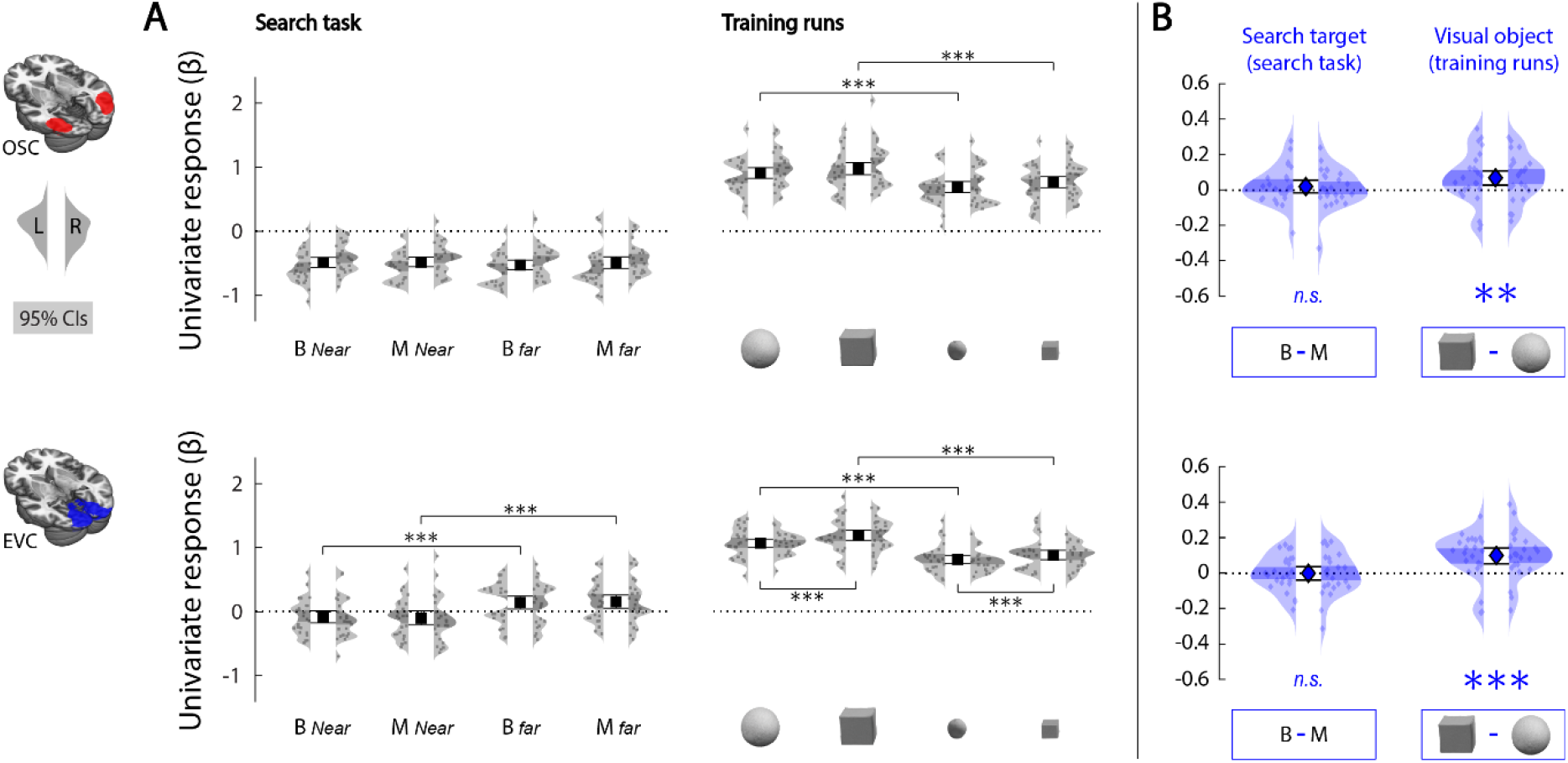
Univariate results. (A) Panel A depicts the average run-based GLM beta-weights for each of the four conditions of the search task (i.e., activity evoked during near and far melon and box search), and the training runs (i.e., activity evoked by viewing large and small images of melons and boxes), averaged over all voxels within a ROI. (B) Panel B depicts univariate contrasts, derived from the values reported in Panel A. In the left plot, positive values reflect a stronger BOLD response evoked following a ‘B’-cue (box) compared to an ‘M’-cue (melon) in the search task. In the right plot, positive values reflect a stronger BOLD response evoked by viewing an image of a box compared to melon in the training runs. To summarize the findings: in the search task, we found no difference in BOLD response during search preparation for melons compared to boxes, in either OSC (p = .190) or EVC (p = .474). In the training runs, images of boxes evoked a stronger BOLD response than images of melons in OSC (p = .004) as well as EVC (p < .0005). Based on this, it appears unlikely that the multivariate cross-classification of search target from visual objects (reported in Figures 2B and 3 of the main manuscript) can be fully accounted for by a shared difference in univariate responses. Small gray or blue dots represent the difference in average beta-weight between pairs of conditions for individual participants, obtained separately from the left and right hemispheres (displayed within the left and right kernel-density plots, respectively). The central markers reflect the population mean, averaged across hemispheres. Error bars around the central markers, and shaded areas within the kernel-density plots represent the bootstrapped 95% confidence intervals of the mean. Asterisks denote significance in one-sample or paired bootstrap tests (2000 samples). \*\**p* < .005, \*\*\**p* < .0005.

**Supplemental Figure S3.**
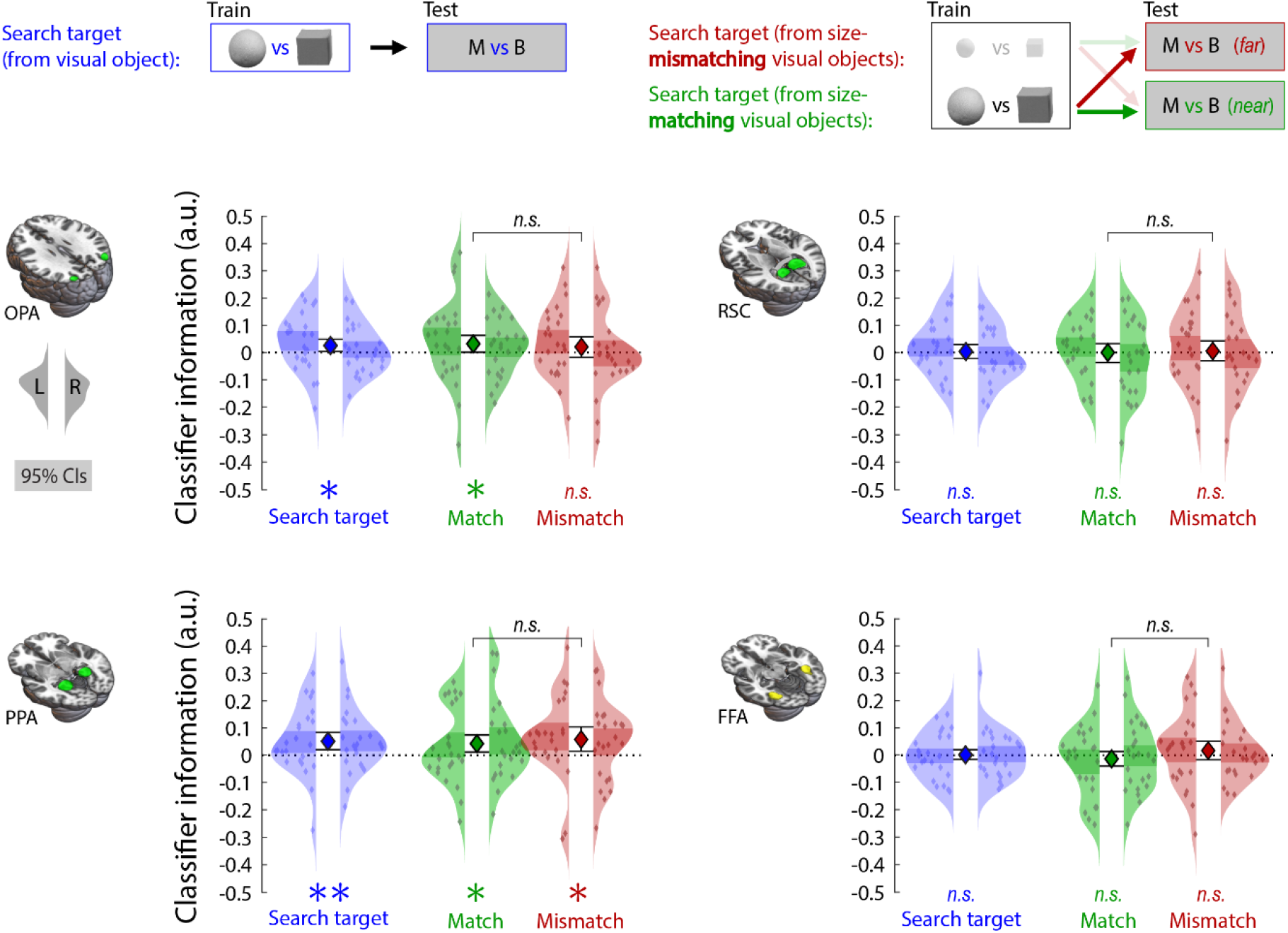
Main analyses applied to scene-selective and face-selective regions. This figure depicts the main (cross-decoding) analyses reported in Figure 2B and 3 of the manuscript for four other regions of interest: scene selective regions OPA (occipital place area), PPA (parahippocampal place area), and RSC (retrosplenial cortex), and face-selective region FFA (fusiform face area). Following the same procedure as that of the object-selective region of interest reported in the main manuscript (OSC), the scene-selective ROIs were obtained by (1) identifying voxels that exhibited a significantly (*p*_uncorrected_ < .05) stronger response to scenes than to objects in the functional localizer runs, and (2) intersecting these scene-selective voxels with three different population-level functionally-defined scene-selective masks (retrieved from ^[S3]^), reflecting OPA, PPA and RSC. Similarly, the face-selective region was obtained by (1) identifying voxels that exhibited a significantly stronger response to faces than to objects and scenes in the functional localizer runs, and (2) intersecting these face-selective voxels with a population-level functionally-defined FFA mask (retrieved ^[S3]^). The results show that two scene-selective regions (OPA and PPA) exhibit an object-specific preparatory bias (depicted in blue). These regions’ preferential responses to rectilinear stimuli and cardinal orientations ^[S4, S5, S6]^ might underly their ability to differentiate between melons and boxes in our study. The ability to distinguish between melons and boxes, however, did not improve when the classifier was trained on objects of the appropriate size as compared to the inappropriate size given the current viewing distance. Small colored dots represent classifier information (derived from distance-to-bound) for individual participants, obtained separately from the left and right hemispheres (displayed within the left and right kernel-density plots, respectively). The central markers reflect the population mean, averaged across hemispheres. Error bars around the central markers, and shaded areas within the kernel-density plots represent the bootstrapped 95% confidence intervals of the mean. \**p* < .05, \*\**p* < .005.

